# Lateral plate mesoderm cell-based organoid system for NK cell regeneration from human pluripotent stem cells

**DOI:** 10.1101/2021.12.03.471069

**Authors:** Dehao Huang, Jianhuan Li, Fangxiao Hu, Qitong Weng, Tongjie Wang, Huan Peng, Bingyan Wu, Chengxiang Xia, Hongling Wu, Jiapin Xiong, Jiali Lin, Yunqing Lin, Yao Wang, Qi Zhang, Xiaofei Liu, Lijuan Liu, Xiujuan Zheng, Haiyan Qiu, Yang Geng, Xin Du, Lei Wang, Jie Hao, Jinyong Wang

## Abstract

Human pluripotent stem cell (hPSC)-induced NK (iNK) cells are a promising “off-the-shelf” cell product for universal immune therapy. Conventional methods for iNK cell regeneration from hPSCs include embryonic body-formation and feeder-based expansion steps, which bring instability, time-consuming, and high costs for manufacture. In this study, we develop an embryonic body-free, organoid aggregate method for NK cell regeneration from hPSCs. In a short time window of 27-day induction, millions of hPSC input can produce over billions of iNK cells without the necessity of NK cell-expansion feeders. The iNK cells highly express classical toxic granule proteins, apoptosis-inducing ligands, as well as abundant activating and inhibitory receptors. Functionally, the iNK cells eradicate human tumor cells by mechanisms of direct cytotoxity, apoptosis, and antibody-dependent cellular cytotoxicity. This study provides a reliable scale-up method for regenerating human NK cells from hPSCs, which promotes the universal availability of NK cell products for immune therapy.

## INTRODUCTION

Immunotherapy provides a new paradigm for saving patient lives with malignancies. Adoptive transfer of natural T cells or NK cells already shows significant therapeutic effects in patients with several tumor types, including CD19^+^ B cell malignancies, breast cancer, and ovarian cancer.^1, 2^ However, immunotherapy still faces problematic issues related to limited cell source, economic costs, and urgent manufacture necessity, as well as low efficiency of gene-editing/engineering.^3–5^ Human pluripotent stem cells (hPSCs), including embryonic stem cells (ESCs) and induced pluripotent stem cells (iPSCs) are ideal to overcome the above-mentioned problems.^4, 6^ Human ESCs, as natural early-stage pluripotent stem cells, have safety advantages over iPSCs due to rare gene mutation accumulations and avoidance of reprogramming process by exogenous genes.^7, 8^

Unlike T cells, NK cells have unique advantages in immune therapy due to little toxicities and no stringent requirement of HLA-matches.^3^ NK cells inherit the power to directly kill those tumor cells that utilize the typical T-cell-escaping mechanism of down-regulating HLA class I (MHC-I) molecules. In the presence of therapeutic monoclonal antibodies, NK cells can further improve their anti-tumor efficacy via antibody-dependent cellular cytotoxicity (ADCC) mechanism.^9^ Human PSC-derived NK cells even showed promising results in eliminating tumor cells *in vivo*.^10, 11^ Hence, regenerative NK cells from hPSCs are considered as an ideal “off-the-shelf” immune cell product for translational medicine.

Encouragingly, hPSCs have been successfully induced into hematopoietic and immune lineage cells *in vitro* via an interim step of EB formation.^12–15^ Hematopoietic lineages originate from the lateral plate mesoderm (LM) during embryonic development.^16^ Conventionally, EB formation and monolayer differentiation methods can both generate mesoderm progenitors from hPSCs with low and variable efficiencies.^17, 18^ Thus, the elevation of the yields of lateral plate mesoderm cells may improve the terminal yields of regenerative immune cells from hPSCs. The OP9 feeder cell line (M-CSF deficient) and derivatives are commonly used for hematopoietic differentiation and lympogenesis *in vitro*.^19^ Fetal thymic organ cultures (FTOCs) show great advantages over monolayer induction system in T cell induction *in vitro*.^20, 21^ Artificial thymic organoid (ATO) culture also promote the T cell regeneration from hPSCs.^22^ Thus, three dimensional structure of stromal cells benefits hemogenesis and lympogenesis.

In this study, we developed an embryonic body-free, organoid aggregate method for NK cell regeneration from human pluripotent stem cells. We firstly optimized the production efficiency of lateral plate mesoderm intermediates, which is essential for the initiation of NK-specific organoid aggregates. Secondly, organoid aggregates are prepared by mixing LM and OP9 stromal cells and were induced toward NK cell development following conventional NK cell induction conditions. In a short time window of 27-day induction, one million hPSC input can produce over one billion iNK cells without the necessity of NK cell-expansion feeders. The iNK cells exhibited strong anti-tumor activity in animal models bearing human tumor cells.

## RESULTS

### Efficient induction of lateral plate mesoderm from hPSCs

A typical phenomenon in hematopoietic induction in petri dishes is that primitive hematopoietic progenitor cells (HPCs) lacking lymphoid-lineage potential are always predominant.^23, 24^ Definitive hematopoietic cell fate commitment in natural embryos undergoes sequential events of lateral plate mesoderm (LM) formation, hemogenic endothelial cells (HECs) specification, endothelial to hematopoietic transition (EHT), and the emergence of definitive hematopoietic progenitors.^16, 25^ To address the low efficiency of NK lymphogenesis, we combined optimizing strategies and established a scheme for regenerating NK cells from hPSCs, which comprises stepwise generations of abundant early-stage LM cells,^26, 27^ HECs, definitive HPCs,^22^ and mature NK cells **(Fig. 1a)**.^28^ Cell images and real-time quantitative PCR results showed that human pluripotent stem cells began to migrate on the first day of differentiation in petri dishes, and highly expressed the primitive streak-specific gene *TBXT* **(Fig. 1b, c)**.^29^ On the second day, the cells proliferated rapidly, and expressed lateral plate mesoderm markers *APLNR* and *HAND1*.^18, 30^ The paraxial mesoderm-specific gene *MSGN1* was not activated during the entire lateral plate mesoderm induction process **(Fig. 1b, c; Supplementary Fig. S1)**.^26^ Flow cytometry analysis also showed that more than 60 percent of the cells on the first day of differentiation expressed the BRACHYURY protein, which is encoded by the *TBXT* gene. On the second day of differentiation, over 90% cells expressed the apelin receptor (APLNR) **(Fig. 1d, e)**. Thus, the lateral plate mesoderm cells are efficiently produced from hPSCs within 48-hour monolayer induction.

**Fig.1.**
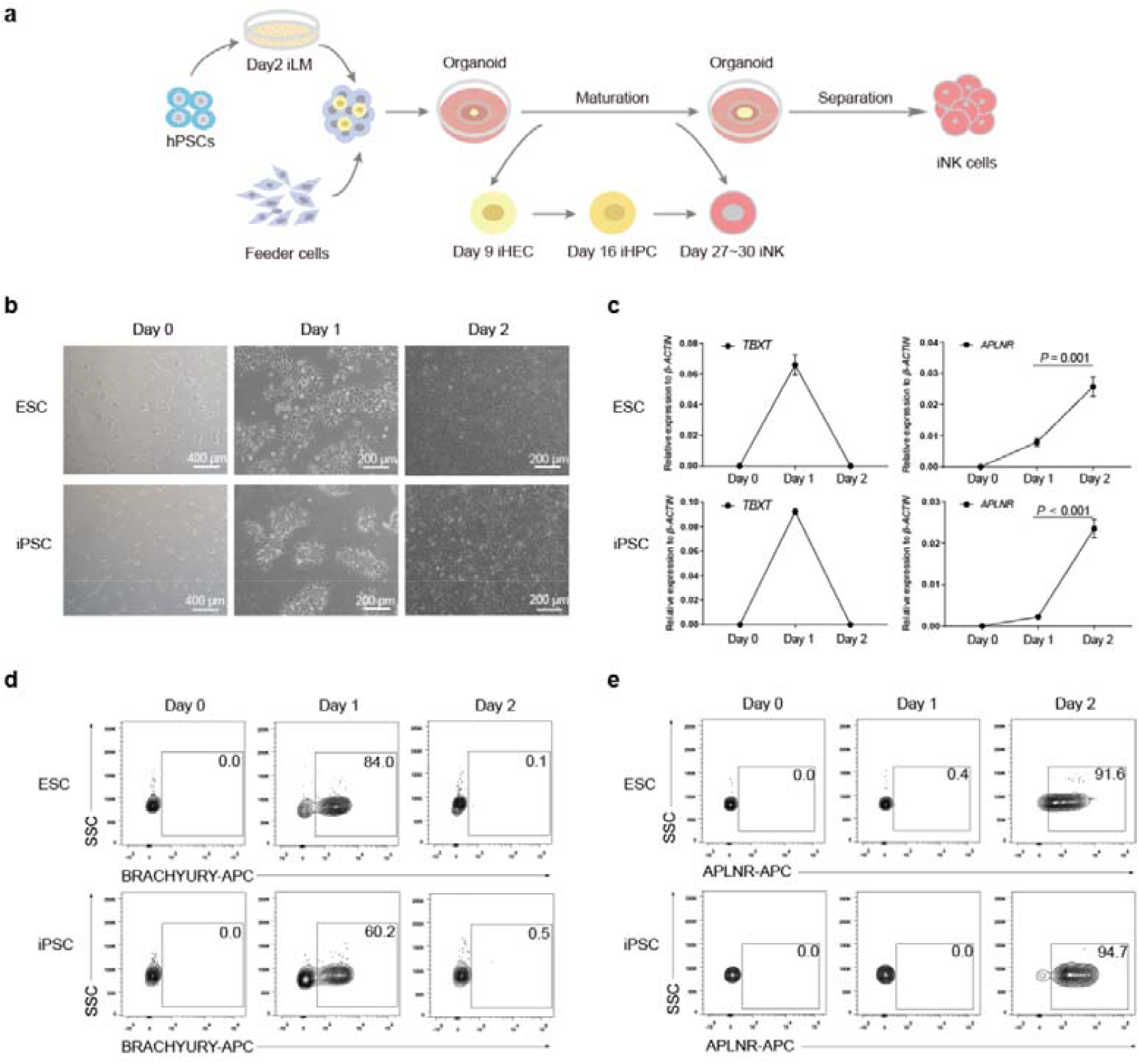
Lateral plate mesoderm induction from human pluripotent stem cells. **a** Schematic diagram of NK cell regeneration. **b** Cell images at specified time points. **c** Real-time quantitative PCR analysis of *TBXT* and *APLNR* gene expression levels at selected time points. β*-ACTIN* gene was used as internal control. The relative gene expression levels were calculated as 2^-ΔCt^. Statistics: two-tailed independent *t*-test. **D** Dynamic analysis of the ratios of BRACHYURY^+^ primitive streak (PS) cells at selected time points by flow cytometry. **e** Dynamic analysis of the ratios of APLNR^+^ lateral plate mesoderm (LM) cells. ESC: human ESC line hPSC-2; iPSC: human iPSC line hPSC-6.

### Organoid aggregate system for NK cell regeneration

We combined the induced lateral plate mesoderm (iLM) and OP9 feeder cells to prepare organoid aggregates at Day 2. The images of organoid aggregates at Day 3, Day 16, and Day 27 were shown **(Fig. 2a)**. Further, we observed the occurrence of abundant induced hemogenic endothelial cells (iHEC, CD144^+^% in CD73^-^DLL4^+^CD31^+^CD34^+^ : >98.5%) on Day 9, and induced hematopoietic progenitor cells (iHPC, CD43^+^% in CD235a^-^CD45^+^CD34^+^ : >98.6%) on Day 16.^31^ On Day 27, over 97.8% cells already showed the terminal mature NK cell phenotypes (iNK, CD45^+^CD3^-^CD56^+^CD16^+/-^) **(Fig. 2b, c, d)**.^32^ Morphologically, ESC-iNK and iPSC-iNK cells were similar to the activated UCB-NK cells **(Fig. 2e)**. Statistically, the organoid aggregates produced 3.54 ± 0.40 (mean ± SD) million CD56^+^ iNK cells at Day 27 and reached 4.10 (median) million at Day 30 **(Fig. 2f)**. We tested as many as six hPSC lines derived from six individuals and all these stem cell lines reproducibly generate abundant iNK cells (*P* <0.001) **(Fig. 2g)**. On average, one

**Fig.2.**
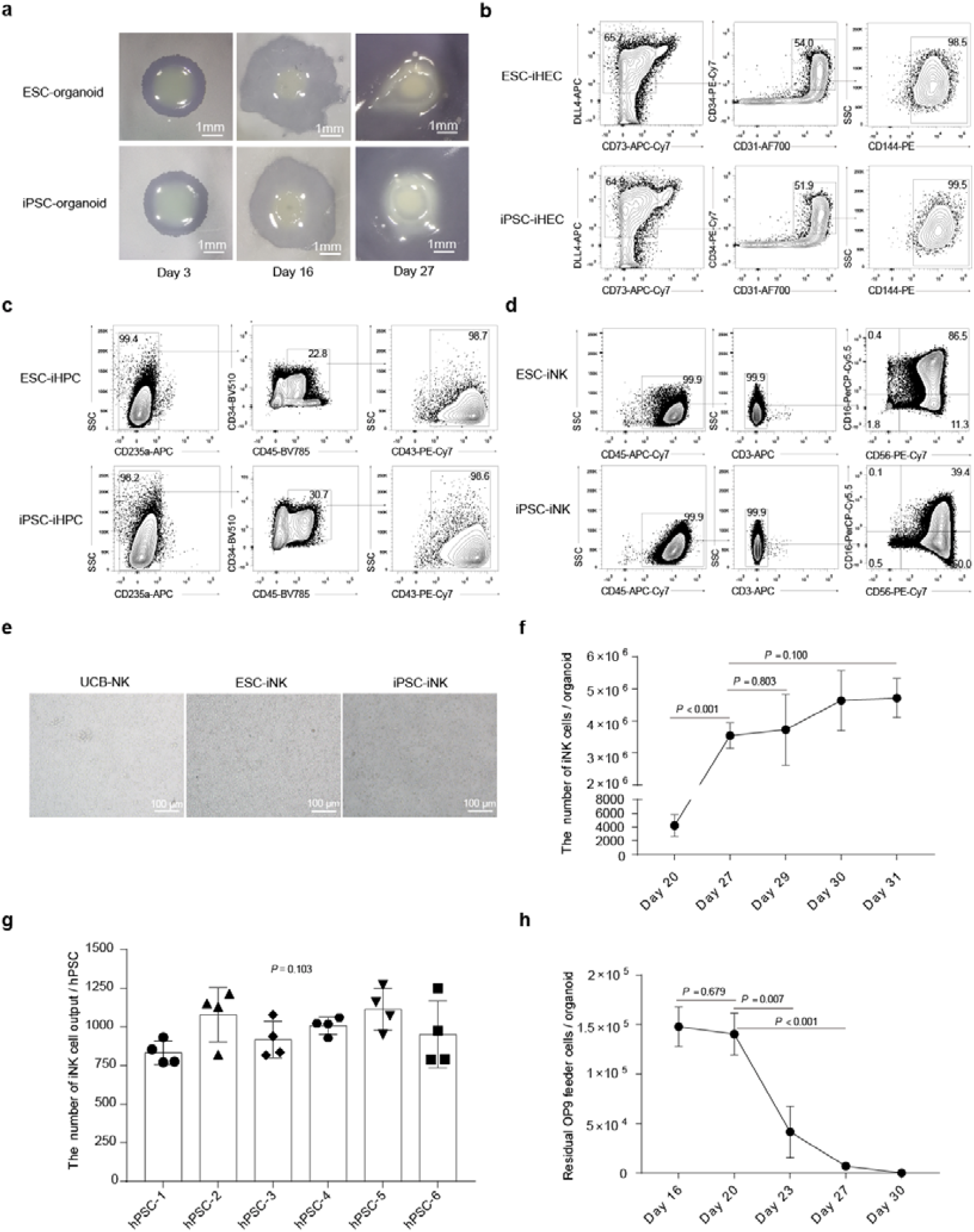
Stepwise induction of iNK cells from lateral plate mesoderm using organoid system. **a** Organoid aggregate images at specified time points. **b** Immuno-phenotypes of hemogenic endothelial cells (iHEC, CD73^-^DLL4^+^CD31^+^CD34^+^CD144^+^) at Day 9. **c** Immuno-phenotypes of hematopoietic progenitor cells (iHPCs, CD235a^-^CD45^+^CD34^+^CD43^+^) at Day 16. **d** Immuno-phenotypes of induced NK cells (CD45^+^CD3^-^CD56^+^CD16^+/-^) at Day 27. **e** The images of NK cells separated from organoid aggregates at Day 27. **f** The number of CD45^+^CD3^-^CD56^+^ iNK cells at the indicated time points (n = 3 each group). Statistics: two-tailed independent *t*-test and Mann-Whitney U test. **g** iNK cell yields per hPSC input. Statistics of iNK cells at Day 27 (n = 4 each group). Data were collected from four batches of induction experiments using five human ESC lines (hPSC-1, hPSC-2, hPSC-3, hPSC-4, and hPSC-5) and one iPSC line (hPSC-6). One point represents the mean of iNK cell output from three repeats per batch induction experiment. Statistics: one-way ANOVA. **h** The residual OP9 feeder cells at the selected time points (n = 3 each group). Statistics: two-tailed independent *t*-test.

hPSC can output 983.56 ± 32.43 (mean ± SD) iNK cells at Day 27 **(Fig. 2g)**. Interestingly, the residual OP9 feeder cells decreased sharply at Day 20 and almost disappeared at Day 30 (*P* <0.001) **(Fig. 2h)**. In addition, we assessed multiple OP9-derived cell lines for organoid aggregates, including OP9-hDLL1, and OP9-hDLL4, which showed comparable iNK cell regeneration efficiencies **(Supplementary Fig. S2)**. Collectively, an organoid aggregate system effectively supports NK cell regeneration starting from LM cells.

### iNK cells express specific activating receptors, inhibitory receptors, and effector molecules

To illustrate the molecular features of iNK cells, we performed single-cell RNA-seq of iNK cells and natural control NK cells from umbilical cord blood (UCB-NK). The expression patterns of activating receptors and inhibitory receptors determine the activating status and functionalities of NK cells.^33, 34^ As expected, both ESC-iNK and iPSC-iNK cells expressed classical activating and inhibitory receptor genes, including activating receptor genes *NCR3, NCR2, KLRK1*, and *SLAMF7*, and inhibitory receptor genes *KLRC1*, and *CD96*.^34, 35^ We also confirmed the expression of these receptors on iNK cells at protein levels by flow cytometry analysis **(Fig. 3a, b)**. The type II transmembrane protein CD94 dimerizes with NKG2A to form CD94/NKG2A heterodimers, which recognize non-classical HLA-E class I molecules and inhibit signal activation or combine with NKG2C to form CD94/NKG2C receptor to transmit activating signals and promote NK cell activation.^36^ Indeed, the CD94 molecule on the surface of iNK cells were very abundant (CD94^+^% in CD56^+^ : >99.7%) **(Fig. 3b)**. In addition, the iNK cells also highly expressed the essential effector molecules, including apoptosis-related ligands (TRAIL and FasL), cytotoxic granules (GzmB and perforin), and activating molecule CD69 **(Fig. 3a, b)**.^34^ Collectively, iNK cells express the typical NK cell markers and effector molecules.

**Fig.3.**
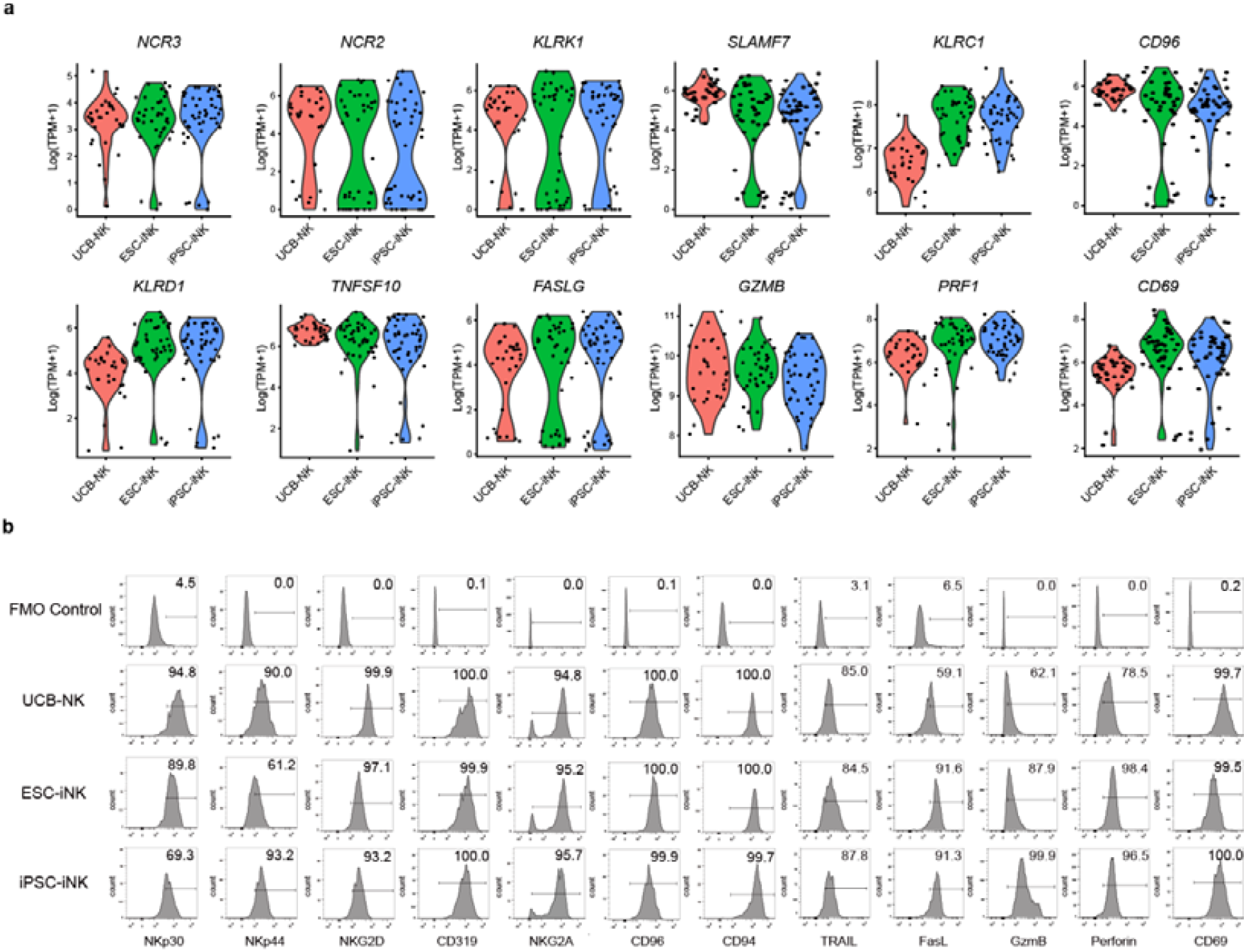
Molecular features of ESC-iNK, iPSC-iNK, and UCB-NK cells. **a** Violin plots show the expression profiles of indicated NK cell-surface receptors and effectors (*NCR3, NCR2, KLRK1, SLAMF7, KLRC1, CD96, KLRD1, TNFSF10, FASLG, GZMB, PRF1*, and *CD69*) in ESC-iNK (n = 41), iPSC-iNK (n = 47), and UCB-NK (n = 32) cells. The expression value (TPM) of each gene was transformed with log2. One point represents one cell. **b** The expression levels of NK cell typical receptors and effectors (NKp30, NKp44, NKG2D, CD319, NKG2A, CD96, CD94, TARIL, FasL, GzmB, Perforin, and CD69) were analyzed by flow cytometry. ESC: human ESC line hPSC-2; iPSC: human iPSC line hPSC-6.

### iNK cells eliminate tumor cells *in vitro* and possess ADCC activity

Unlike adaptive T cells, NK cells can directly recognize and kill tumor cells with downregulated HLA class I (MHC-I) expression.^37^ Therefore, we chose a NK cell-sensitive human ovarian cancer cell line A1847 (tdTomato^+^) for validation of the tumor-killing ability of iNK cells. ESC-iNK, iPSC-iNK and UCB-NK cells at sequential doses (5 ×10^3^, 1 ×10^4^, 2 ×10^4^, and 5 ×10^4^) were respectively co-cultured with 1 ×10^4^ A1847-tdTomato^+^ tumor cells. As expected, all three sources of NK cells can efficiently target A1847 cells and lead to immediate apoptosis of tumor cells within 4 hours **(Fig. 4a, b)**. Consequently, the viable numbers of A1847-tdTomato^+^ cells sharply decreased after 4-hour coculture with iNK cells at various E:T ratios **(Fig. 4c)**. NK cells can enhance tumor-killing effect by a mechanism of antibody-dependent cellular cytotoxicity (ADCC).^38^ In order to examine the ADCC ability of iNK cells, we combined the anti-CD20 antibody (Rituximab) with iNK cells for killing CD20^+^ human lymphoma cell line Raji at different E:T ratios (1:1, 2:1, 5:1, 10:1, 20:1).^39^ As expected, the anti-CD20 antibody further enhanced the iNK cell killing activity against Raji tumor cells **(Fig. 4d, e, f)**. Taken together, iNK cells derived from hPSCs possess tumor-killing activity via direct-and ADCC-manners.

**Fig.4.**
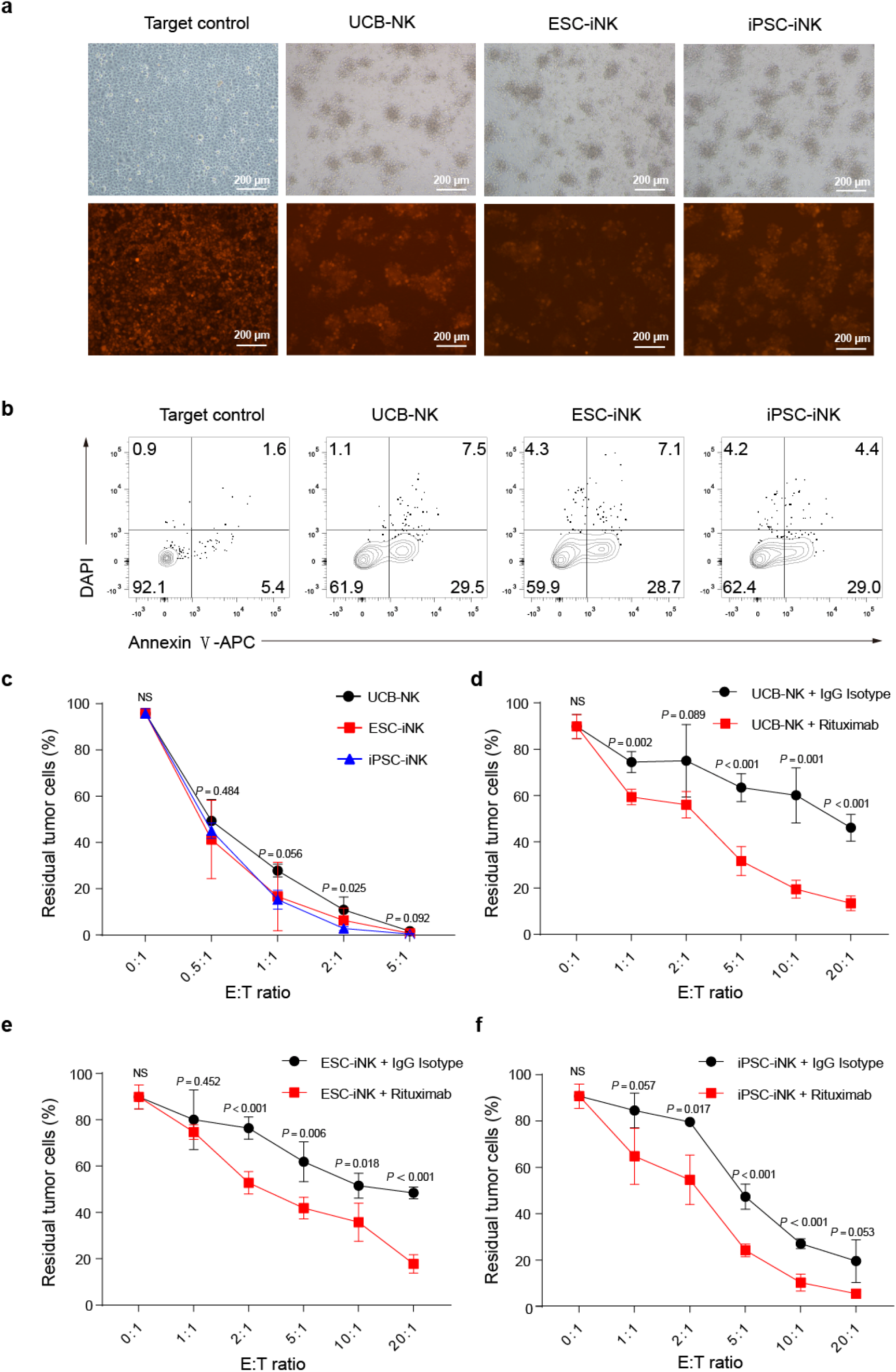
iNK cells exhibit tumor-killing activities *in vitro*. **a** Representative images of residual A1847-tdTomato^+^ tumor cells after 4-hour coculture with iNK cells (E:T = 5 : 1 ratio). **b** Flow cytometry analysis of apoptotic A1847-tdTomato^+^ tumor cells after 1 hour co-culture with NK cells. Gated populations indicate the status of tumor cells: necrotic cells (DAPI^+^ Annexin V^-^), late apoptotic cells (DAPI^+^ Annexin V^+^), early apoptotic cells (DAPI^-^ Annexin V^+^), and living cells (DAPI^-^ Annexin V^-^). **c** Ratios of residual A1847-tdTomato^+^ tumor cells were measured after 4-hour coculture. 1 ×10^4^ target (T) cells (A1847-tdTomato^+^ tumor cells) were cocultured with effector (E) cells (ESC-iNK cells, iPSC-iNK cells, or UCB-NK cells) at 5 ×10^3^, 1 ×10^4^, 2 ×10^4^, and 5 ×10^4^ doses respectively (n = 6 each group). Data were collected from two independent experiments. Statistics: one-way ANOVA and Kruskal-Wallis test. **d-f** Raji cells were incubated with NK cells by adding human IgG1 isotype control antibody (20 μg/mL) or anti-CD20 antibody (Rituximab, 20 μg/mL) for 4 hours. Ratios of residual Raji tumor cells were measured after 4-hour coculture with ESC-iNK, iPSC-iNK or UCB-NK cells. A total of 2 ×10^3^ Raji tumor cells as target (T) cells were cocultured with NK cells (2 ×10^3^, 4 ×10^3^, 1 ×10^4^, 2 ×10^4^ and 4 ×10^4^) as effector (E) cells for 4 hours at different E:T ratios (1:1, 2:1, 5:1, 10:1, 20:1) (n = 4 each group). Data were analyzed by two-sided independent *t*-test. NS, not significant. ESC: human ESC line hPSC-2; iPSC: human iPSC line hPSC-6.

### iNK cells eradicate human tumor cells in xenograft animals

To evaluate therapeutic potential of the iNK cells *in vivo*, we established a human tumor-xenograft animal model by transplanting the luciferase-expressing A1847 (A1847-luci^+^) cells (2 ×10^5^ cells/mouse) in NCG (a commercial NOD/ShiLtJGpt-Prkdc^em26Cd52^Il2rg^em26Cd22^/Gpt strain) mice at Day −1. Then, the iNK cells were intraperitoneally injected into the tumor-bearing animals (1∼1.5×10^7^ cells/mouse) at Day 0 and Day 7. Meanwhile, UCB-NK cells were used as natural NK cell treatment control. IL-2 (10000 IU/mouse) was administrated every two days until Day 21 to enhance the maintenance of human iNK cells in animals.^11^ Bioluminescent imaging (BLI) was performed weekly to signal the kinetics of tumor growth **(Fig. 5a)**. Indeed, both ESC-iNK cells and iPSC-iNK cells efficiently eliminated tumor cells *in vivo* and exhibited comparable tumor-killing efficacies with natural UCB-NK control group. Meanwhile, the tumor burden of the tumor only group became more and more severe, as indicated by the radiance and the value of total flux (p/s) **(Fig. 5b, c)**, which eventually needed euthanasia due to heavy tumor burden between Day 37 and Day 41 post A1847-luci^+^ cells injection. The ESC-iNK cell treated group and iPSC-iNK cell treated group survived significantly longer than the tumor-only control mice (Tumor only: 38 days, Tumor + UCB-NK: > 87 days, Tumor + iPSC-iNK: > 82 days, Tumor + ESC-iNK: > 120 days, p < 0001) **(Fig. 5d)**. Collectively, these results show that the ESC-or iPSC-derived iNK cells can efficiently kill tumor cells and prolong survivals of tumor-bearing animals.

**Fig.5.**
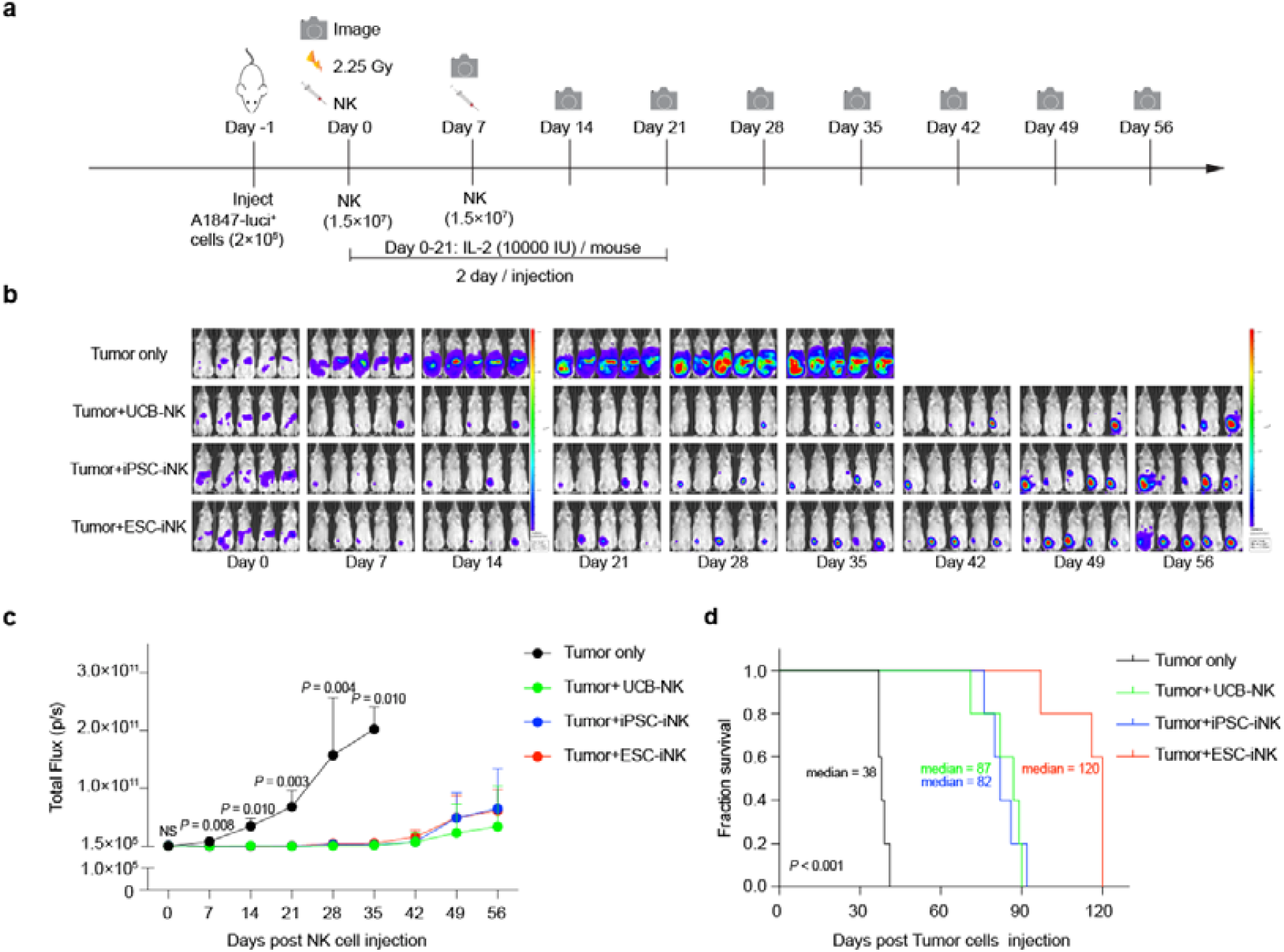
ESC- or iPSC-derived iNK cells eradicate human tumor cells in xenograft animals. **a** Procedure design for NK cells treatment in tumor-bearing animals. **B** Bioluminescent imaging (BLI) images of the xenograft models (Tumor only, Tumor + UCB-NK, Tumor + iPSC-iNK and Tumor + ESC-iNK, n = 5 each group). The radiance indicates tumor burden. **c** Statistics of the total flux (p/s) of the xenograft models (Tumor only, Tumor + UCB-NK, Tumor + iPSC-iNK and Tumor + ESC-iNK, n = 5 each group). Statistics: Mann-Whitney U test. NS, not significant. **d** Kaplan-Meier survival curves of the xenograft models (Tumor only, Tumor + UCB-NK, Tumor + iPSC-iNK and Tumor + ESC-iNK, n = 5 each group) (*P* <0.001, Log-rank test). Median survival was shown. ESC: human ESC line hPSC-2; iPSC: human iPSC line hPSC-6.

## DISCUSSION

In this study, we develop an embryonic body-free, organoid aggregate method for NK cell regeneration from human pluripotent stem cells. In a short time window of 27-day induction, one million hPSC input can produce over one billion iNK cells without the necessity of NK cell-expansion feeders. Functionally, the iNK cells exhibited strong anti-tumor activity in animal models bearing human tumor cells. Molecularly, the iNK cells showed the typical expression patterns of activating and inhibitory receptors, cytotoxic molecules, and apoptosis-related ligands of natural NK cell counterparts. Mechanistically, the iNK cells possess promising abilities of killing human tumor cells via cytotoxic, ADCC, and apoptotic mechanisms.

In general, cord blood NK cells from one individual can expand up to billions of NK cells following two-week expansion with a conventional expansion protocol in the presence of expanding-feeder cells.^40^ In our study, one batch of hPSCs can regenerate as many as hundred billions iNK cells with the same genetic background and one single hPSC can produce over 1,000 iNK cells in the absence of expansion feeder cells. The iNK cell yields per hPSC in our study are much higher than the conventional EB-induction method (30-100 iNK cells per hPSC).^41^ The efficiency improvements are associated with the generation of pure lateral plate mesoderm intermediates and organoid aggregate incubation approach.

We found that the iNK cells exhibited an activated status from as early as newborn, which is different from natural adult NK cells born with naïve status. This phenomenon raises a possibility that the iNK cells possess less functional specification potential than the diversified natural NK cells developed and distributed in multiple organs in humans.^42^ In our method, the entire NK induction process only takes four weeks, which is much shorter than the time spent by conventional methods.^11, 14^ Regarding the unlimited source of hPSCs, the yields of iNK cells can reach trillions by increasing the input of initiating stem cells. For example, we tested that two million hPSCs can output over 2 × 10^9^ iNK cells within one month induction. For scale up manufacture purpose, one billion hPSCs input can produce over one trillion iNK cells, which might meet the application requirements of 100-1000 patients.^43^

In this study, the iNK cells regenerated from ESC significantly prolonged the survivals of tumor-bearing mice when compared with the groups treated with UCB-NK or iPSC-iNK cells. This phenomenon reappeared even using iNK cells from multiple independent induction assays using ESC as starting cells. It deserves further investigation whether this is the inherited trait of ESC-derived iNK cells or is caused by different genetic backgrounds of starting cell lines. In future, more genetic background strains of hPSCs needs to be broadly assessed.

In addition, we also found that the proportion of CD16^+^ iNK cells derived from different stem cells fluctuates with a range from 30 to 90%. However, the rates of CD16^+^ iNK cells from the same stem cell line are very stable. Thus, the causing factor of CD16 expression variations is associated with different hPSC lines rather than the induction method itself **(Supplementary Fig. S3a, b)**. This observation also brings an advantage to pre-screen master stem cell lines capable of producing iNK cells with high expression rates of CD16, as CD16^+^ iNK cells is crucial for combinational application of targeting antibodies in treating cancer patients.

In conclusion, we develop an embryonic body-free, organoid aggregate method for NK cell regeneration from human pluripotent stem cells. One million hPSC input can produce over one billion iNK cells without the necessity of NK cell-expansion feeders within one month induction procedure. The iNK cells are able to kill tumor cells via cytotoxic, apoptosis-inducing and ADCC mechanisms. Our study provides a guide for manufacturing large-scale “off-the-shelf” NK cell products for immune therapy.

## MATERIALS AND METHODS

### Cell culture

Human ESC lines (hPSC-1, hPSC-2, hPSC-3, hPSC-4, and hPSC-5) were provided by National Stem Cell Resource Center, Institute of Zoology, Chinese Academy of Sciences. Human iPSC line (hPSC-6) was derived from a healthy donor’ urine cells by reprogramming, with the donor’ informed consents. All the hPSC lines were maintained in ncTarget medium (Nuwacell) on Matrigel (Corning) coated plates. Umbilical cord blood samples were obtained from Guangdong Cord Blood Bank (Guangzhou, China) and the UCB-NK cells were cultured in the KBM581 medium (Corning) supplemented with IL-2 (200 UI/mL, Miltenyi) and SGR-SM (1%, DAKEWE). OP9 cell line was purchased from ATCC and cultured with α-MEM (Gibco) with 20% fetal bovine serum (FBS) (Ausbian). A1847 cell line was purchased from Honsun Biological Technology Co., Ltd (Shanghai, China) and cultured in RPMI 1640 Medium (Gibco) supplemented with 10% FBS (Ausbian). Raji cell line was purchased from the Cell Resource Center of Shanghai Institutes for Biological Sciences, Chinese Academy of Sciences (Shanghai, China) and cultured in RPMI 1640 Medium (Gibco) supplemented with 10% FBS (Ausbian).

### Hematopoietic differentiation and NK cell regeneration *in vitro*

Briefly, for lateral plate mesoderm differentiation, the Accutase (400-600 units/mL, Sigma-Aldrich)-digested hPSCs were resuspended in the ncTarget medium (Nuwacell) supplemented with thiazovivin (0.5 μM, Selleck), and then were plated on the vitronectin (10 μg/mL) coated dishes. Particularly, lateral plate mesoderm differentiation medium was the TeSR™-E6 basal medium (STEMCELL Technologies) supplemented with BMP4 (40 ng/mL, R&D Systems), ACTIVIN A (30 ng/mL, PeproTech), bFGF (20 ng/mL, PeproTech), CHIR99021 (6 μM, Selleck) and PIK-90 (100 nM, Selleck) at Day 0, and supplemented with BMP4 (40 ng/mL, R&D Systems), A-83-01 (1 μM, Selleck) and C59 (1 μM, Selleck) at Day 1. After 24-hour induction, lateral plate mesoderm cells were digested with Accutase and 0.5 mM EDTA solution (1:1) for 1 min and resuspended in hematopoietic differentiation medium (HDM) which components were TeSR™-E6 medium supplemented with 10 μM SB431542 (selleck), 10 μM Hydrocortisone (selleck), 5 ng/mL Flt3L (PeproTech), 5 ng/mL TPO (PeproTech), 50 ng/mL SCF (PeproTech), 50 ng/mL EGF (PeproTech), 50 ng/mL VEGF (R&D System), 50 ng/mL bFGF (PeproTech), 50 ng/mL IGF-1 (PeproTech), 50 μg/mL ascorbic acid and 2% SGR-SM (DAKEWE). Meanwhile, OP9 feeder cells were harvested by 0.25% trypsinization (Hyclone) and resuspended in HDM. Then 5 × 10^5^ OP9 feeder cells were combined with 2 × 10^4^ lateral plate mesoderm cells to form organoid aggregates.^22^ The organoids were plated on a 0.4 μm Millicell transwell insert (Corning) and placed in 6-well plates containing 1 mL HDM per well. Medium was changed completely every 2-3 days for two weeks. At Day 16, the medium was changed to NK differentiation medium.^28^ Half medium exchanges were performed every 2 days. At Day 27, the mature NK cells were harvested and cultured in the KBM581 medium (Corning) supplemented with IL-2 (200 UI/mL, Miltenyi) and SGR-SM (1%, DAKEWE).

### Real-time quantitative PCR

The total RNA was extracted by the RNAprep pure Cell Kit (TIANGEN), and 1 μg RNA was reversely transcribed into cDNA by using ReverTra Ace® qPCR RT Master Mix with gDNA Remover (TOYOBO). Real-time quantitative PCR was performed with Hieff® qPCR SYBR® Green Master Mix (Yeasen Biotechnology). β*-ACTIN* was used for normalization. The relative gene expression level was calculated as 2^-ΔCt^. All primers used in this study were listed in the **Table 1**.

### Flow cytometry

Organoid was incubated in 1 mL digestion buffer (1 mg/mL Collagenase type IV (BasalMedia Technologies), 1 mg/mL Dispase (Gibco) and 50U DNase I (Sigma-Aldrich) for 10 minutes at 37°C, mechanically disrupted by pipetting, and incubated again for 10 minutes at 37°C. After complete disaggregation by pipetting, single cell suspensions were prepared by passage through a 70 μm filter (Corning). Expect for organoid, other single-cell suspensions were prepared by Accutase and filtered by 70 μm filter. Cells were blocked by Human TruStain FcX™ (Biolegend, 422302) antibody, and then stained with related antibodies. The following antibodies were used: BRACHYURY (R&D System, IC2085A), APLNR (R&D System, FAB8561A-025), CD3 (Biolegend, HIT3a), CD16 (Biolegend, 302028), CD31 (Biolegend, WM59), CD34 (Biolegend, 343528), CD43 (BD Biosciences, 563522), CD45 (Biolegend, HI30), CD56 (Biolegend, HCD56), CD73 (eBioscience, AD2), CD144 (Biolegend, BV9), CD235a (Biolegend, 349114), DLL4 (Miltenyi, REA1065), NKp30 (Biolegend, P30-15), NKp44 (Biolegend, P44-8), NKG2D (Biolegend, 1D11), CD319 (Biolegend, 162.1), NKG2A (Biolegend, S19004C), CD96 (Biolegend, NK92.39), CD94 (BD Biosciences, HP-3D9), CD69 (Biolegend, FN50), TRAIL (Biolegend, RIK-2), FasL (Biolegend, NOK-1), GzmB (Biolegend, QA18A28), Perforin (Biolegend, dG9). The cells were resuspended in the DAPI (Sigma-Aldrich) solution and were analyzed with BD LSRFortessa X-20 cytometer (BD Biosciences). Flow cytometry data were analyzed by the FlowJo software (Three Star, Ashland OR).

### RNA-seq and data analysis

The cDNA of single UCB-NK, human ESC-iNK and iPSC-iNK cell sorted at Day 27 was generated and amplified using Discover-sc WTA Kit V2 (Vazyme). The quality of amplified cDNA was assessed by qPCR analysis of *GADPH* gene. Samples that passed quality control were used for sequencing library preparation using TruePrep DNA Library Prep Kit V2 (Vazyme). All libraries were sequenced by illumina sequencer NextSeq 500. The raw data (fastq files) were generated using bcl2fastq software (version 2.16.10) and uploaded to the Genome Sequence Archive public database (HRA001609). The raw reads were aligned to human genome hg19 by HISAT2 (version 2.1), and the expression levels in TPM were estimated by StringTie (version 1.3.4).

### Function analysis of hPSCs-derived iNK cells *in vitro*

iNK cells were derived from hPSCs and UCB-NK cells were isolated from umbilical cord blood. UCB-NK cells were stimulated and expanded on IL-21 natural killer cell amplification system (Hangzhou Zhongying Biomedical Technology Co., Ltd) for 14 days. ESC-iNK cells, iPSC-iNK cells and UCB-NK cells(effector, E) were incubated with 1 ×10^4^ A1847-tdTomato^+^ cells (target, T) in Flat-bottom 96-well plates for 4 hours at respective E:T ratios (E:T=0.5:1, 1:1, 2:1, 5:1). For antibody-dependent cellular cytotoxicity (ADCC) analysis, ESC-iNK cells, iPSC-iNK cells and UCB-NK cells (effector, E) were incubated with 2 ×10^3^ Raji-tdTomato^+^ cells (target, T) in U-bottom 96-well plates for 4 hours at respective E:T ratios (E:T=1:1, 2:1, 5:1, 10:1, 20:1). The residual number and apoptotic state analysis of A1847 and Raji tumor cells (tdTmato^+^) were enumerated using the CountBright Absolute Counting Beads (Thermo Fisher), Annexin □ (Biolegend) and DAPI (Sigma-Aldrich) solution by BD LSRFortessa X-20 cytometer (BD Biosciences). Flow cytometry data were analyzed by the FlowJo software (Three Star, Ashland OR).

### Construction of the ovarian cancer xenograft models and treated with iNK cells

NCG mice (NOD/ShiLtJGpt-Prkdc^em26Cd52^Il2rg^em26Cd22^/Gpt, GemPharmatech Co., Ltd.) were intraperitoneally injected with the luciferase-expressing A1847 (A1847-luci^+^) cells (2 × 10^5^ cells/mouse) to construct the ovarian cancer xenograft models at Day −1. Bioluminescent imaging (BLI) (IVIS Spectrum PerkinElmer) was performed on these models to quantitative the tumor burden, and the models with similar total flux (p/s) were randomly divided into four groups (Tumor only, Tumor + UCB-NK, Tumor + iPSC-iNK and Tumor + ESC-iNK) at Day 0. These models were first irradiated (2.25 Gy, Rad Source RS2000), and then intraperitoneally injected with the ESC-iNK, iPSC-iNK, or UCB-NK cells (1-1.5 × 10^7^ cells/mouse) 4 hours post-irradiation and 7 days after the first injection. IL-2 (10000 IU/mouse) was administrated every two days until Day 21 post NK cell injection. BLI was performed every week to trace the tumor cells. Mice suffered from heavy tumor burdens were euthanized for ethical consideration.

### Ethical statement

NCG mice were housed in the SPF-grade animal facility of the Guangzhou Institutes of Biomedicine and Health, Chinese Academy of Sciences (GIBH, CAS, China) and the Institute of Zoology, Chinese Academy of Sciences (IOZ, CAS, China). All procedures of this study were approved by the Institutional Animal Care and Use Committee of the Institute of Zoology, and the Institutional Animal Care and Use Committee of the Guangzhou Institutes of Biomedicine and Health. hPSC differentiation toward hematopoietic and immunology lineage cells, and related anti-tumor activity assessments of iNK cells in animals are approved by the Biomedical Research Ethics Committee of the Institute of Zoology, Chinese Academy of Sciences. The iPSC preparations from donor somatic cells and related assays in this study are also permitted with donor consents.

### Statistics

All quantitative analyses were based on at least three sample replicates. Two-tailed independent *t*-test, Mann-Whitney U test, one-way ANOVA, Kruskal-Wallis test and Kaplan-Meier survival curve were performed in the environment of SPSS and GraphPad Prism. Mean values ± SD were shown. Multiple comparisons were determined by one-way ANOVA or Kruskal-Wallis test. Survival curve was calculated using Kaplan-Meier methods.

## ACKNOWLEDGEMENTS

This work was supported by the National Key R&D Program of China (2020YFA0112404), the Strategic Priority Research program of the Chinese Academy of Sciences (XDA16010601), the National Key R&D Program of China (2019YFA0110203), and the National Natural Science Foundation of China (81925002, 32100904). We thank Tiancheng Zhou, and Tian Zhang from Dr. Guangjin Pan’s lab for guiding iPSC preparation and culture.

## AUTHOR CONTRIBUTIONS

D.H., JH.L., and F.H. designed and conducted experiments, performed data analysis, and wrote the manuscript. Q.W. and Y.L. performed RNA-Seq and data analysis. T.W., H.P., B.W., C.X., H.W., J.X., JL.L., Y.W., Q.Z., X.L., L.L., X.Z., H.Q, Y.G., L.W., and J.H. participated in multiple experiments; J.W., D.H., J.L., and F.H. wrote the manuscript; X.D. discussed the manuscript, and J.W. designed the project and provided the final approval of the manuscript.

## Figure legends

**Supplementary Fig. S1.**
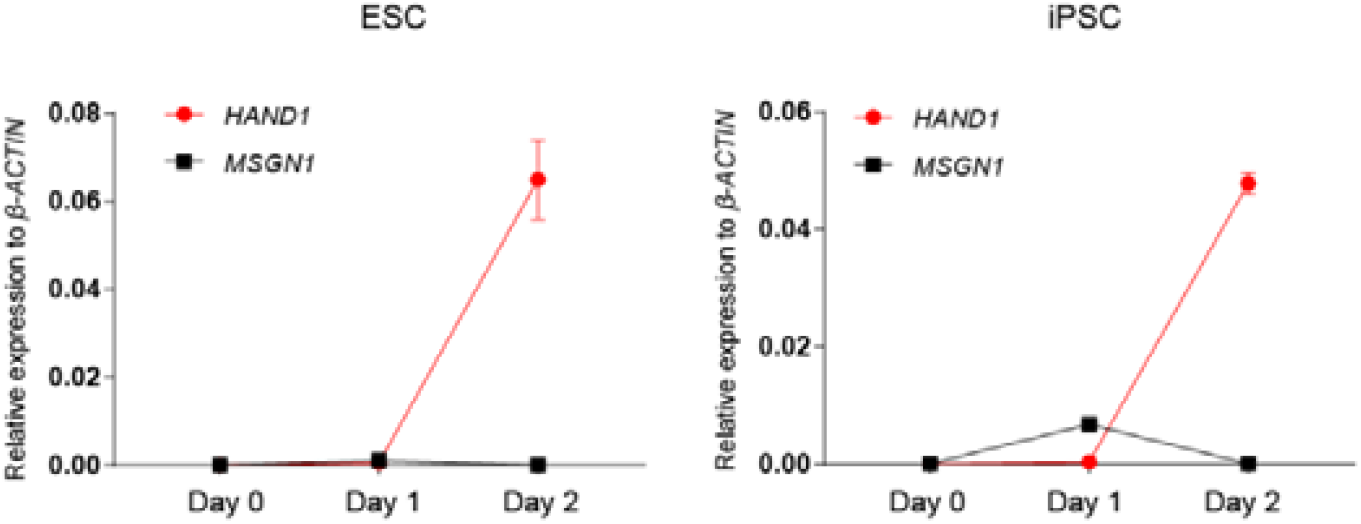
Real-time quantitative PCR analysis of *HAND1* and *MSGN1* gene expression levels. Real-time quantitative PCR analysis of *HAND1* and *MSGN1* gene expression levels at Day 0, Day 1, and Day 2. ESC: human ESC line hPSC-2; iPSC: human iPSC line hPSC-6.

**Supplementary Fig. S2.**
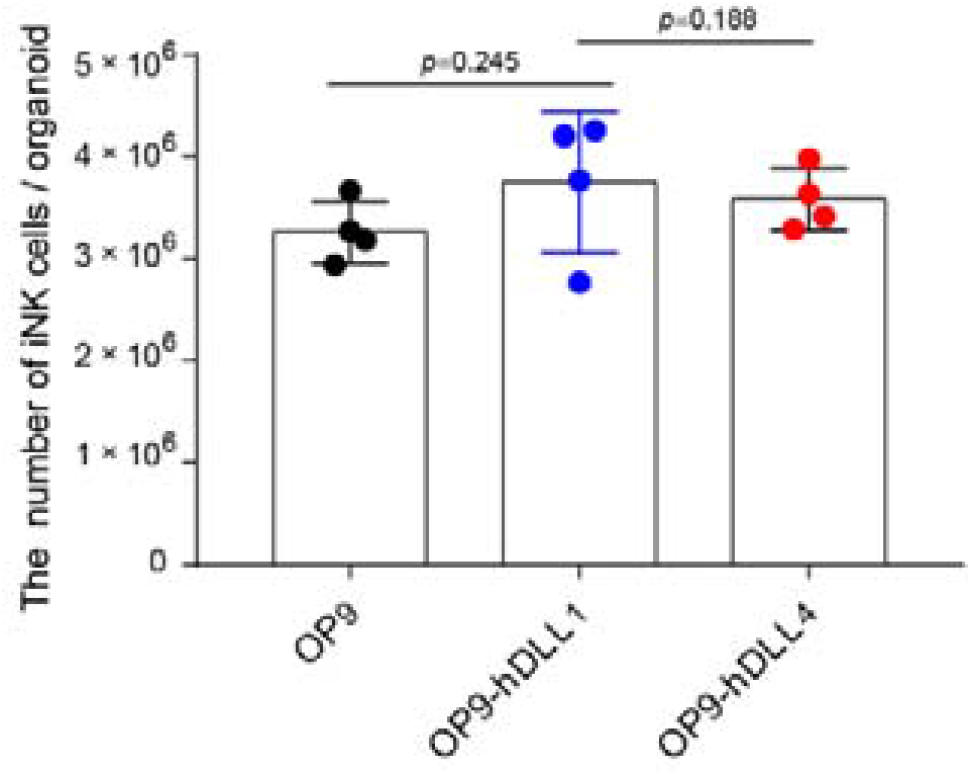
Comparison of NK cell productions from organoid aggregates composed of OP9, OP9-hDLL1, and OP9-hDLL. 5 ×10^5^ OP9, OP9-hDLL1, and OP9-hDLL4 feeder cells were combined with 2 ×10^4^ mesoderm cells separately to form organoid aggregates at Day 2. Data were collected and analyzed at Day 27 (n = 4 each group). Statists: two-tailed independent *t*-test.

**Supplementary Fig. S3.**
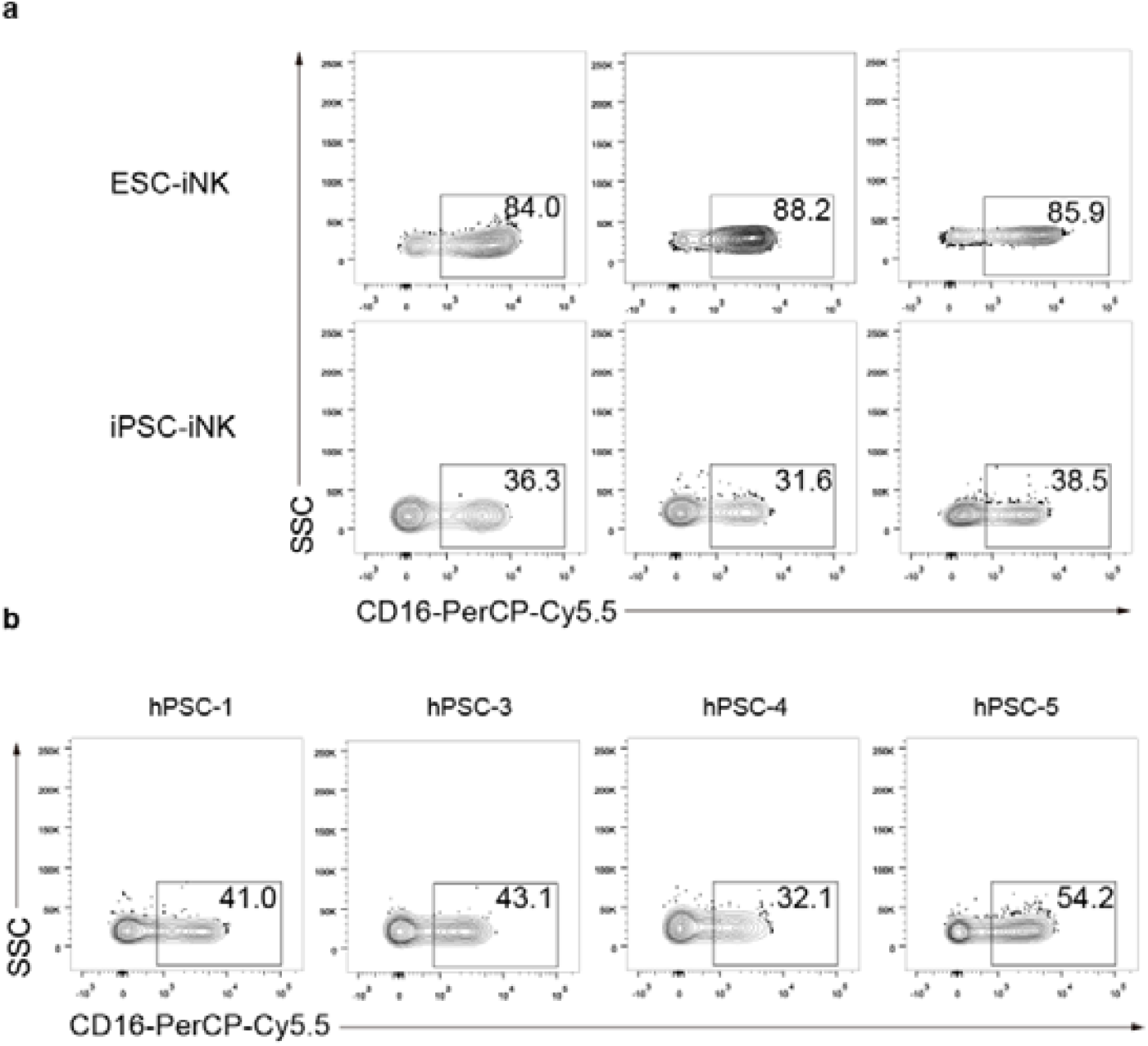
CD16 molecule expression pattern of iNK cells. **a** CD16 expression level of human ESC line hPSC-2-derived iNK cells and human iPSC line hPSC-6-derived iNK cells. Data were collected from three independent experiments. CD16^+^ cells were gated on CD45^+^CD3^-^CD56^+^ iNK cells. **b** CD16 expression level of human embryonic cell lines hPSC-1, hPSC-3, hPSC-4, and hPSC-5-derived iNK cells. CD16^+^cells were gated on CD45^+^CD3^-^CD56^+^ iNK cells. All samples were collected at Day 27.

**Supplementary Table S1.**
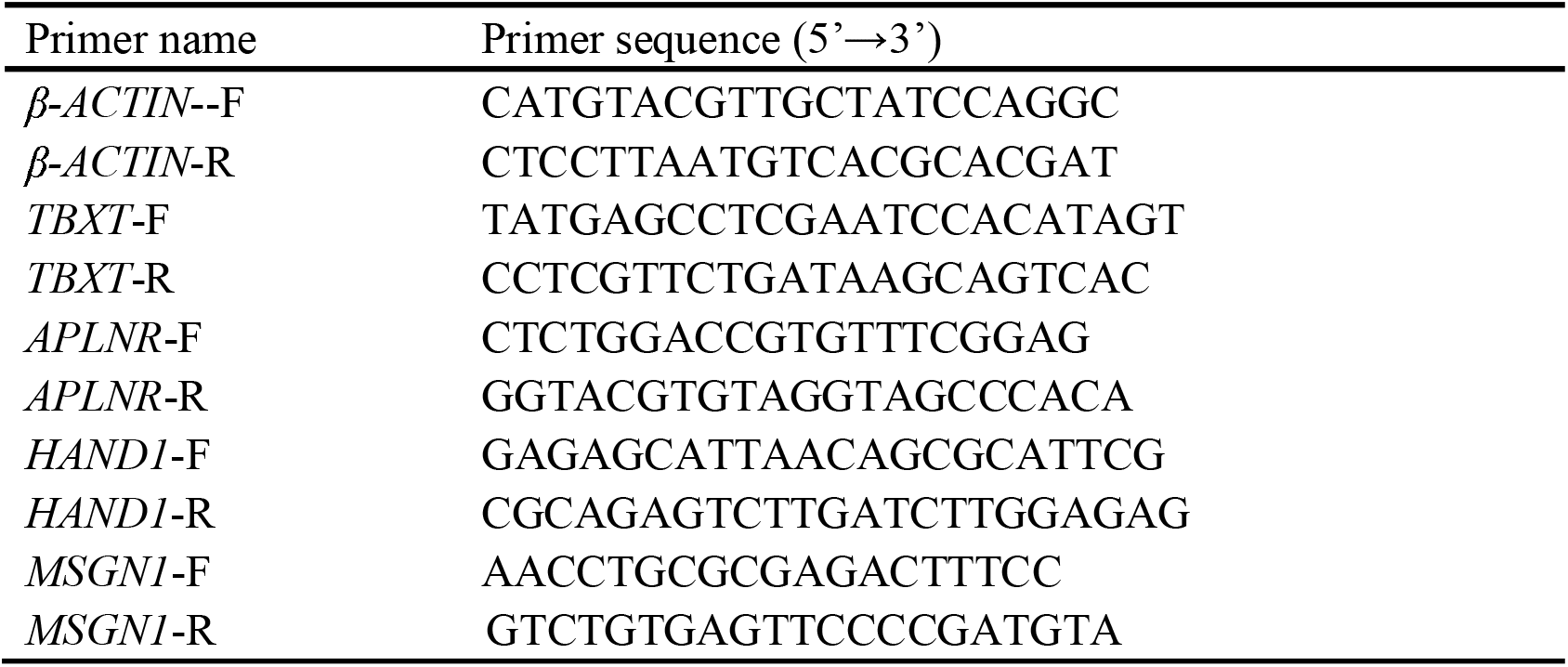
Primer sequences for real-time quantitative PCR of selected genes

